# Protein massively parallel binding assay reveals transcription factor binding determinants

**DOI:** 10.1101/2025.03.05.641616

**Authors:** Offir Lupo, Sagie Brodsky, Tamar Jana Lang, Wajd Manadre, Gili Valinsky, Yoav Navon, Vladimir Mindel, Naama Barkai

**Author notes:** co-first authors (Offir Lupo, Sagie Brodsky, Tamar Jana Lang).

## Abstract

Transcription factors select their genomic binding sites in genomes depending on their DNA binding domain (DBD) but also on regions outside the DBD (nonDBD). However, it remains challenging to define these determinants within nonDBDs and reveal their mechanism of action. Towards this, we introduce here an *in-vivo* method for parallel analysis of thousands of designed peptides for binding a DNA sequence of interest (Protein Massively Parallel Binding Assay, pMPBA). We apply it to scan the full sequence space of budding yeast TFs and generate a detailed map of DNA localizing determinants. Within the set of predicted DBDs, we reveal a large variation in DNA binding affinities, depending on the family and on different sequence characteristics, including charge. Strong signals were not confined to predicted DBDs but included a considerable fraction of nonDBD peptides, most of which were predicted as intrinsically disordered. pMPBA opens new possibilities for high-throughput analysis of peptide-DNA binding within cells.

## Introduction

Transcription factors (TFs) regulate gene expression by binding to specific gene regulatory regions in genomes. DNA binding domains (DBDs) within TFs localize to short DNA sequence motifs that are present within bound regulatory regions. However, these short DBD-recognized motifs are insufficient for predicting the TF genomic preferences since most genomic occurrences remain unbound^1–3^. It is therefore widely recognized that TF regions outside the DBD (nonDBD) contribute to the selection of TF target sites. Defining respective nonDBD sequences directing genomic binding, however, remains challenging^4–10^.

DBDs bind their specific motif through base-specific interactions, mostly within the major groove^11^. These interactions are enabled by DBD folding into DNA-compatible structure, and multiple non-specific interactions made with the DNA backbone^11^. DBDs are therefore classified to a limited set of conserved structural families which are readily predicted by their amino acid sequence. By contrast, regions outside the DBD can influence DNA binding through different means including interactions with co-binding TFs, association with histone marks, or recognition of non-canonical DNA folds. These regions lack signatures and, in general, are difficult to predict through sequence analysis.

Complicating the analysis of TF nonDBDs is their enrichment with intrinsically disordered regions (IDRs)^12–19^. Studies of sequence-function relations within IDRs lag these of globular domains, mostly due to the rapid divergence of IDR sequences^20–22^, at least when tested using the common, alignment-based comparative methods. New sequence comparative methods that search for conserved features within IDR sequence are emerging^21–28^, but these are not yet of sufficient resolution to predict TF regions directing genomic binding. Limiting method development is the scarcity of experimental data defining functional IDR sequences, which is needed for directing and refining sequence-based predictions.

Towards understanding the sequence-function relation of TF IDRs, we recently interrogated dozens of budding yeast TFs through systematic nonDBD truncations, and further used extensive sets of designed mutations that spread across the hundreds of residues composing two model IDRs^1,29–34^. The effect of each of these mutations on binding preference was defined by profiling genomic binding, revealing that most TFs contain multiple specificity-conferring regions that are spread across their sequence.

In this paper, we adapt a complementary approach of scanning the TF sequence space for short peptides that individually localize to DNA within cells. This peptide scanning borrows from recent studies that scanned TFs for activation domains (ADs), namely peptides that induced reporter gene expression when fused to a DBD of choice^35–41^. However, as we were interested in DNA-binding or localization, rather than transcription induction, we needed a new method that will report on peptide-DNA association, independent of gene transcription.

To achieve this, we modified our recently published method, the Massively Parallel Binding Assay (MBPA)^42^ which measures binding of a given TF of interest to thousands of DNA sequence inside cells, to work in the complementary direction, namely measuring binding of thousands of designed peptides to a DNA sequence of interest. We denote this new method protein Massive Parallel Binding Assay (pMPBA). As its main advantage, pMPBA provides a transcription-independent readout of the instantaneous peptide-DNA association through a rapid library-based high throughput analysis. We apply this method to scan the full sequence space of budding yeast TFs for peptides that bind a long non-specific DNA region. Our results reveal significant variation in DNA binding across predicted DBDs and identified strong DNA localization by peptides outside the DBD, most of which characterized by high sequence disorder. We discuss the implications of these results for our understanding of the TF-target search process in genomes.

## Results

### Protein Massively Parallel Binding Assay for defining DNA interacting protein domains

To enable parallel analysis of DNA binding by thousands of designed peptides, we extended the MPBA (Massive Parallel Binding Assay) approach we recently introduced^42^. MPBA measures binding of thousands of designed DNA sequences to a given protein of interest. We now extended it to compare the binding of thousands of designed peptides to a DNA sequence of interest.

In pMPBA, the tested protein-DNA pairs are integrated into plasmids and transformed into cells. The system is engineered such that the binding of the plasmid-carrying peptide to the tested DNA results in cleavage of the plasmid. In its current implementation, plasmid cleavage is measured within the region that includes the coding sequence of the tested peptides and can therefore be quantified by sequencing. To achieve this, pMPBA is implemented as follows: First, a plasmid is engineered to contain the tested DNA, adjacent to a library integration site (**Figure 1A**). Second, a library of peptides is designed and cloned into the plasmid, ensuring that each plasmid carries a single variant. Critically, within the plasmid, the peptide is fused to an MNase and a nuclear localization signal and is expressed under a strong promoter (TEF1). As an additional feature, we linked each peptide to a URA3 selection marker using the ribosome-skipping inducer (E2A) sequence^43^, ensuring that only in-frame peptides are retained. Third, the plasmids are transformed into cells, and DNA cleavage is triggered by MNase activation through a short calcium pulse^44^. Finally, PCR amplification of a region covering the cleaved site and the peptide sequence is performed and quantified by high-throughput sequencing. The temporal changes in the abundance of each peptide sequence, relative to cells in which MNase was not activated, providing an estimate of the relative DNA occupancy compared to all tested peptides.

**Figure 1.**
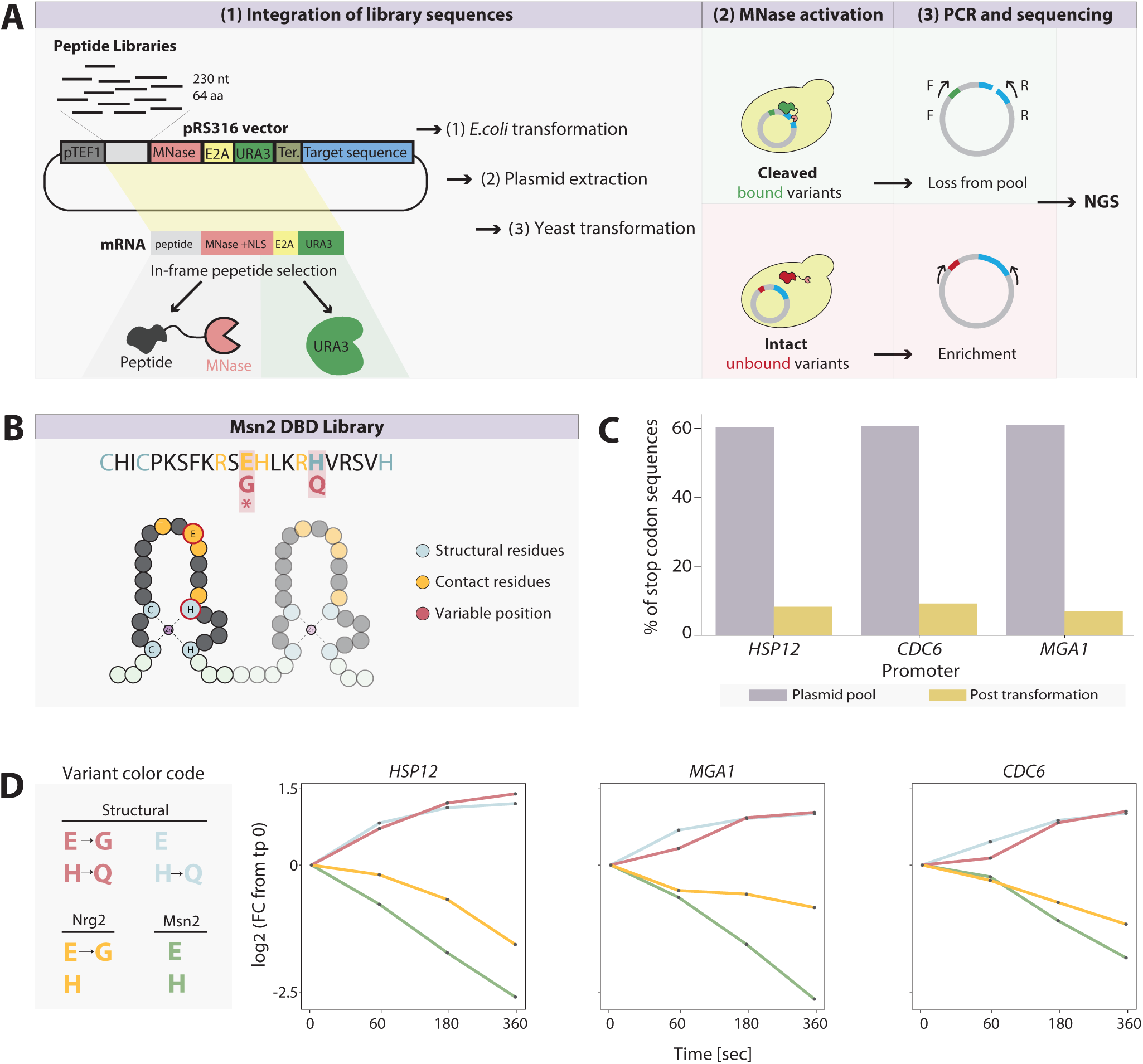
Peptide Massive parallel binding assay (pMPBA) using library of short peptides. **(A)** *pMPBA for peptide variants workflow*: A target sequence of interest is inserted into the plasmid, then, a library of sequences coding for 64AA peptides is integrated upstream to an MNase **(1, top)**. Plasmid pool is then transformed into bacteria for propagation followed by plasmid extraction. A ribosome skipping inducing sequence (E2A) is located between the peptide-MNase and a URA3 gene **(1, bottom)**. Following yeast transformation, only cells expressing an in-frame peptide will grow in uracil depleted media. A short calcium pulse activates the MNase, triggering the cleavage of peptide-bound plasmids **(2)**. Next, a targeted PCR is performed, amplifying non-cleaved sequences covering the region between the peptide-coding sequence to the target promoter sequence. Amplicon sequencing before and after MNase activation then provides a quantitative binding measurement for each variant **(3)**. **(B-D)** *Method validation*: To test our system, we chose the zinc-finger DBD of the Msn2 TF. **(B)** We modified its first zinc-finger domain to generate a few variants: (1) Premature stop codon (E->*) (2) DNA contact residue modification to match its paralog Nrg2 (E->G), (3) Structural mutation predicted to disrupt proper 3D folding of the domain (H->Q), and (4) Combination of mutation (2) and (3). This library was integrated into three plasmids, each containing a different target promoter (HSP12, CDC6, and MGA1). **(C)** Ribosome skipping peptide reduces incorrectly translated proteins: The fraction of library sequences containing a premature stop codon is shown before (grey) and after (orange) yeast transformation. **(D)** pMPBA successfully quantifies variant binding: Library binding was tested over an MNase activation time course. The log2 fold-change from time-point zero (no MNase activation) is shown for the three tested promoters.

Note that peptide binding can be tested across any DNA of interest, provided that the tested region is linked to the encoded peptides through e.g., a peptide-linked barcode. Here, we are interested in a broad range of DNA-bound peptides, and therefore considered a long DNA region that included the full peptide-coding sequence (230 bps), as well as an MNase, NLS, E2A, URA3 gene (1744 bps), and a selected promoter sequence which we varied between experiments (∼540-750 bps), totaling ∼ 2600 bps.

We validated our approach using a small peptide library of DBD mutants. This library included the native DBD of the Msn2 TF, and three additional variants: one inserting a premature stop codon, one replacing an essential structural residue to glutamine (H660Q) and one replacing a DNA-contacting residue to that of a related TF, Nrg2 (E655G, **Figure 1B**). Applying pMPBA to measure library binding to three DNA sequences differing in the promoter region confirmed the expected results: variants including the premature stop-codon were eliminated following yeast transformation, validating the efficiency of the E2A selection for in-frame peptides (**Figure 1C)**. Further, plasmid carrying the WT DBD were depleted from the pool following MNase activation, with levels decreasing by ∼6-fold relative to the DBD structural mutant at 360 second activation (**Figure 1D)**. Lastly, the contact residue mutant showed a partial decrease in signal, consistent with binding at reduced affinity. Of note, these measured occupancies were highly similar between the three tested promoters, indicating that the binding signal is dominated by non-specific association with the long DNA sequence rather than specific promoter binding.

### DBD-annotated domains show a range of DNA binding capacities

Across genomes, DBDs of conserved families are predicted through sequence-based annotations, but only a fraction of these predicted DBDs were tested experimentally. As a first application of pMPBA, we compared binding of annotated DBDs within a set of 136 budding yeast TFs. For this, we collected the full set of domains predicted within the associated sequences by common annotation databases (pfam^45^, SMART^46^ or SUPERFAMILY^47^), totaling after refinement in 198 unique peptides. Of these, 167 corresponded to annotated DBDs while the remaining, annotated to different functions and served as controls.

We synthesized the respective domain library and tested its binding to nine DNA sequences differing by the tested promoters **(Figure 2B)**. Changes in library composition were reproducible between repeats (r=0.86) and mostly independent of the tested promoter (r=0.89), as above, indicating non-specific association with the large DNA sequence tested. We used the median depletion across all promotes as our initial measure of binding signal.

**Figure 2.**
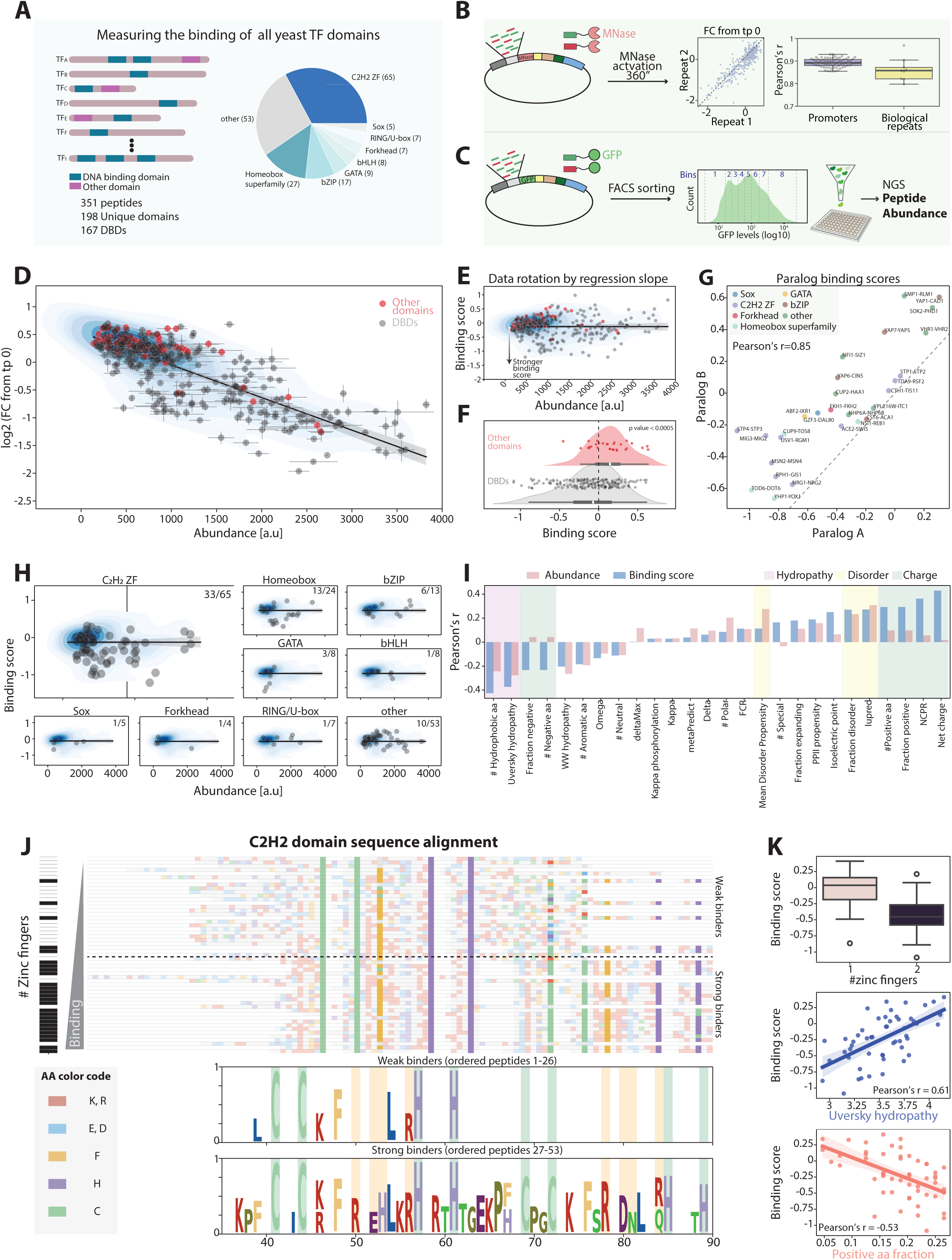
Comprehensive analysis of yeast transcription factor DNA-binding domains. **(A)** *Yeast TF domain library, a scheme*: A library was created using all pfam^45^, SMART^46^, and SUPERFAMILY^47^ annotated domains. Out of 351 total sequences, 198 originated from unique domains (methods), encompassing a total of 167 different TF DBDs. The fraction of each DBD family in the library is shown. **(B-C)** *Abundance and binding quantification:* The library was integrated into nine plasmids, each containing a different target promoter (Table S5) and tested for binding after 360 seconds of MNase activation **(B, left)**. Peptide fold-change from time 0 (no activation) showed high reproducibility between repeats and target promoters **(B, right-top)**. The same library was also integrated into a plasmid containing a GFP reporter gene instead of an MNase **(C)**. Following transformation and ∼20 growth cycles (methods), the cells were sorted into 8 bins based on fluorescence, grown and sequenced to measure domain abundance. **(D-E)** *Peptide binding correlates with abundance*: The log2 fold-change from time-point zero of each domain is shown as a function of protein abundance quantified using GFP. Data density is shown in contour. Error bars on the x-axis represent the two GFP repeat measurements, and on the y-axis, the two biological repeats after averaging the signal over all promoters (methods)**. (E)** Shown is the abundance-normalized binding score as a function of peptide abundance. To calculate binding score, a linear regression between log2 fold change from time 0 and abundance was calculated. Then, the distance of each point from the regression line was computed and referred to as binding score (methods). For visualization, data rotation according to the regression slope was performed. **(F)** *Strong binding of DBDs compared to other TF domains:* The distributions of binding scores for TF DBDs (grey) and other TF domains (red) are shown. Each dot represents a unique domain. The p-value of a two-sample T-test is shown. **(G)** *DBDs of close paralogs demonstrate highly similar binding strength:* The binding scores for DBDs belonging to TF ohnologs (Whole genome duplication retained paralogs) are shown. Pairs are colored based on the DBD family, as indicated. List of ohnologs was taken from^68^ **(H)** *Strong binding demonstrated by Zinc finger DBDs:* For each DBD family, the binding score is shown as a function of peptide abundance (as in **E**). The number of domains exhibiting strong binding (lower than regression line) is indicated for each TF family. **(I)** *Positive charge positively correlates with binding score:* The binding scores and abundance of all domains were correlated with multiple peptide sequence properties^51,56,65^. Note the high correlation of net and positive charge with binding score but not with abundance. Note the positive correlation of predicted disorder on peptide abundance and the negative correlation of hydrophobic residues on binding score. **(J-K)** *Zinc finger DBD binding is dictated by positive AA fraction and hydropathy:* **(J)** All zinc-finger peptides were aligned based on their consensus sequence (C-2-5x-C-9-12x-H-2-6x-H, where x corresponds to any aa). The aligned amino acid sequences are ordered by binding score, with strong binders found at the bottom. On the left, the number of zinc-finger domains for each measured peptide is shown. The consensus sequences of the weakest and strongest binders are displayed at the bottom, with contact residues highlighted in orange and structural residues in green. **(K)** A boxplot **(top)** shows the distribution of binding scores for peptides containing one or two zinc-finger domains. The zinc-finger DBD binding scores are plotted as a function of hydrophobicity **(K, middle)** and positive amino acid fraction **(K, bottom)**

Binding signals depend on peptide-DNA affinity, but also on peptide abundance. As peptides might be degraded at different rates, we measured the abundance of each peptide in our library using GFP fusion, followed by fluorescence-activated cell sorting (FACS) and sequencing **(Figure 2C and S1A**). Peptides varied in abundance, with highly expressing peptides being mostly of low hydrophobic content and high disorder propensity (**Figure S1B**), consistent with the enrichment of strong degrons with hydrophobic amino acids^48–50^. As expected, binding signals were well correlated with peptide abundance, but still showed significant variation among similarly expressed peptides, indicating differential DNA association (**Figure 2D**). Using this data, we defined peptide binding scores as the abundance-normalized binding measure, obtained by linear regression. This measure is intuitively visualized by rotating the data in the binding-abundance plane (**Figure 2E**). Of note, as we quantify sequence depletion compared to no activation, negative binding scores indicate increased DNA association.

Binding scores were uniformly low for the control domains lacking DBD annotations and were variable across annotated DBDs (**Figure 2F**). Consistent with our present assay capturing mostly non-specific DNA association, differences in DBD binding scores did not correlate with the number of available motifs (**Figure S2A**) and differed between same-family DBDs of same, or similar motif preference (**Figure S2B**). Notably, binding scores were correlated amongst DBD ohnologs (r = 0.85, **Figure 2G**) and strong binders diverged more slowly in evolution, indicating their functional relevance (**Figure S2C**).

The fraction of DBDs that failed to bind DNA in our assay (binding score above the regression line) differed between families. In the largest C2H2 zinc-finger family, 51% (33/65) DBDs were bound, but this fraction was lower in the small families that are, perhaps, less well defined **(Figure 2H; S2D)**. To better understand this difference between bound and unbound DBDs, we searched for distinguishing sequence features^51^ (**Figure 3I)**. Peptides of high hydrophobic content were of generally low abundance and showed poor (abundance-normalized) binding scores. By contrast, peptide charge had no apparent effect on abundance, but was correlated with binding score (Pearson’s r = 0.43)

**Figure 3.**
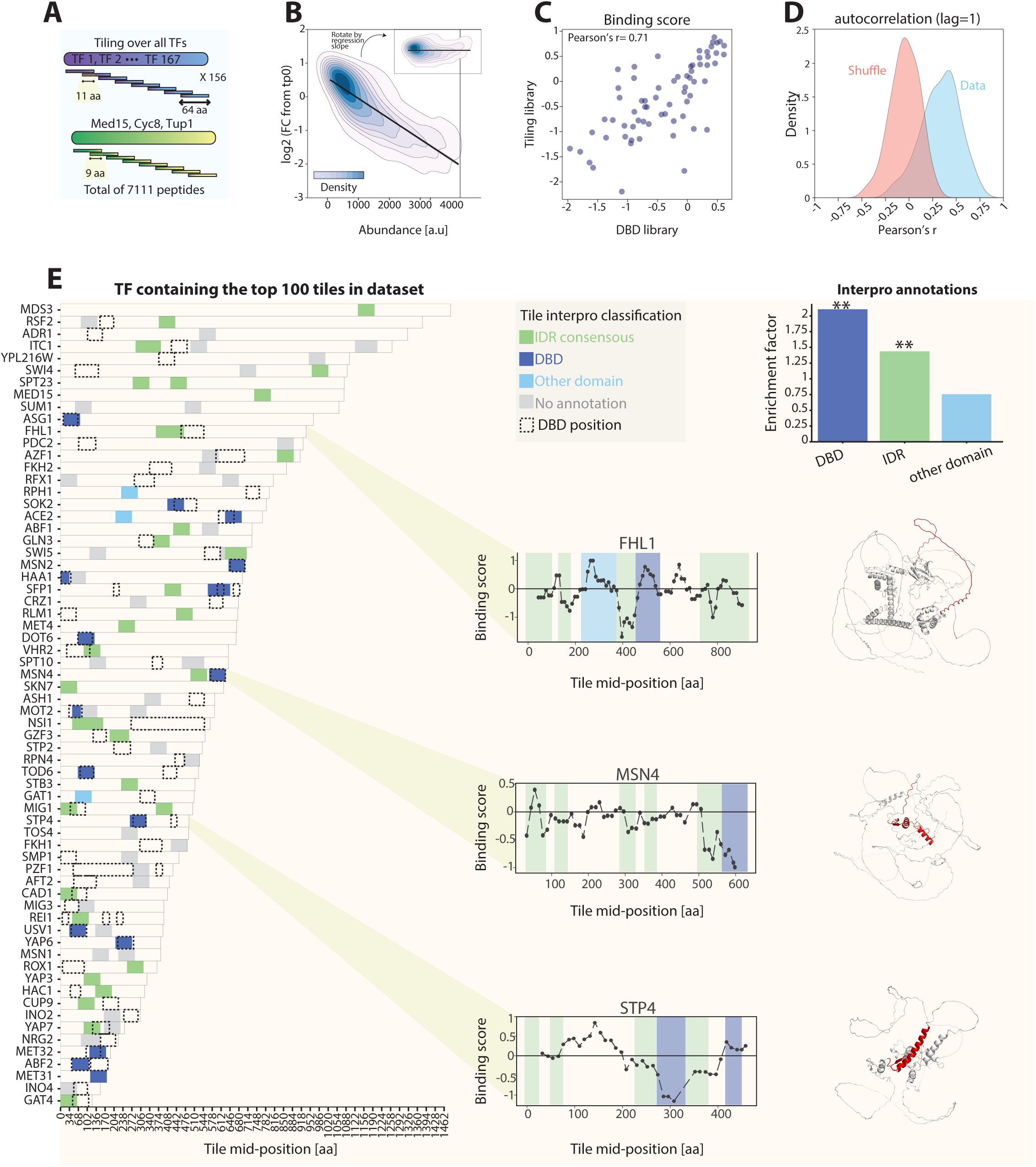
Strong binders are enriched with TF IDR peptides. **(A-B)** *Peptide MPBA for All Yeast TFs:* A tiling library was generated including 153 specific yeast transcription factors (TFs) and three general co-activators **(A, Table S3)**. The library was inserted into multiple plasmids containing different target promoters, and pMPBA was applied as described above. **(B)** The binding log2 fold-change from time point zero (no activation) as a function of peptide abundance is shown as a density contour plot. Date rotation according to the regression slope is also shown. **(C-D)** *High correspondence of TF domain and tiling libraries:* The binding scores of each peptide in the DBD library were compared to those of the closest physical peptide in the TF tiling library **(C)**. The autocorrelation (lag=1) of the log2 fold change from time point zero was calculated for each TF and is shown as a distribution in **(D, blue)**. The peptide order of each TF was shuffled, and the autocorrelation was recalculated and shown as a control **(D, pink)**. **(E)** *Top binders are enriched with DBDs and IDRs:* Shown are the TFs containing the top 100 binding peptides, ordered by amino acid length (left). The DBD of each TF is marked in a black dashed line, and the top binding peptides are colored based on their InterPro^52^ annotation (methods) as indicated. The enrichment scores of the indicated groups are shown as bars **(top right)**. The binding scores and the AlphaFold^54,55^ predicted structure shown for three example TFs. Regions with an average binding score lower than −0.7 are colored in red.

To verify these correlations of peptide binding scores with charge and hydrophobicity, we considered same-family DBDs, focusing on the C2H2 zinc finger family. Aligning all domains based on their defining features revealed stronger binding of domains containing two adjacent zinc-fingers (**Figure 2J**). As seen in the full DBD set, also here binding scores correlated most strongly with charge and low hydropathy (**Figure 2K**). We conclude that pMPBA distinguishes functional DBDs of general DNA affinity, with the strongest binders enriched in positive charge and low hydrophobicity.

### NonDBD-derived peptides display a range of DNA binding comparable to annotated DBDs

We next applied pMPBA to scan the full TF sequences for peptides that associate with the tested DNA. For this, we generated a tiling library of the 136 TFs above, adding 17 TFs not previously included. In this design, we scanned protein sequences at a 11 aa resolution, with sequential peptides sharing 53 of their 64 residues. We further added three general co-factors scanned at higher resolution (Med15, Cyc8, and Tup1, at 9-aa spacing), reaching a total of 7111 peptides (**Figure 3A**). Binding was tested against five DNA regions differing in the selected promoter, all of which again led to similar changes in peptide depletion patterns (median Pearson’s r =0.58 and r=0.64 between promoters and repeats, respectively, **Figure S3A**). We next measured peptide abundance as above and defined an abundance-normalized binding score (**Figure 3B**). This GFP measure displayed higher variance within the more complex library (**Figure S3B**), and we therefore limited our analysis to 4421 peptides showing a reliable GFP measure (methods).

The TF tiling library included peptides that overlapped these analyzed in the domain library above. Comparing binding scores of these overlapping (although not identical) peptides showed the expected high correlation (r=0.71, **Figure 3C**). Using a tiling design further enabled to test reproducibility within each library, as overlapping sequences are more likely to have similar binding signal. This was indeed seen in an autocorrelation analysis, further confirming the reproducibility of our assay (**Figure 3D**).

Next, we focused on the 100 top-bound peptides (**Figure 3E, S3C)**. As expected, these peptides were enriched with annotated DBDs. However, DBD-overlapping peptides were still a minority (17/100) in this set of top-scoring peptides. Most of the remaining high-scoring peptides were not annotated within the databases we initially used, and we therefore examined the broader InterPro database^52^. In addition to DBD annotation, the “IDR consensus regions”^53^ annotation was assigned to 35 of the 100 top-bound peptides. AlphaFold predictions^54,55^ were consistent with these annotations, as can be seen for two representative examples of high-scoring peptides: A disordered region within the Fhl1 TF separating the DBD and the SMAD domain, and a disordered region adjacent to the Msn4 DBD. For comparison, STP4, for which only the DBD scored highly is also presented. More generally, IDR consensus regions found among the 100 top binders were located at different regions within the TFs, both in proximity but also further away from the annotated DBDs.

### TF non-DBD-derived binding determinants are disordered and enriched in positive charge

A large fraction of the 100 top-scoring peptides were therefore annotated as “consensus IDR”. To test this association more systematically, we asked whether peptides located in disordered regions are more likely to be associated with the tested DNA than peptides located to structural domains lacking DBD annotations. For this, we separated the sequence of each TF into three regions, corresponding to the DBD, and to the disordered or structural regions outside the DBD. Indeed, the proportion of bound peptides in both DBDs and disordered non-DBDs were comparable and were twice as high as these within structured non-DBD regions (41%, 47% and 21%, respectively, **Figure 4A**). Exemplifying this is Cyc8 which lacks annotated DBD but shows high binding scores exclusively at peptides located to the 50% of its sequence annotated as disordered (**Figure 4B**-**C**). Notably, while TF IDRs often include activation domains (ADs), we found no correlation between binding scores and AD-signatures^38^ (**Figure S4A**).

**Figure 4.**
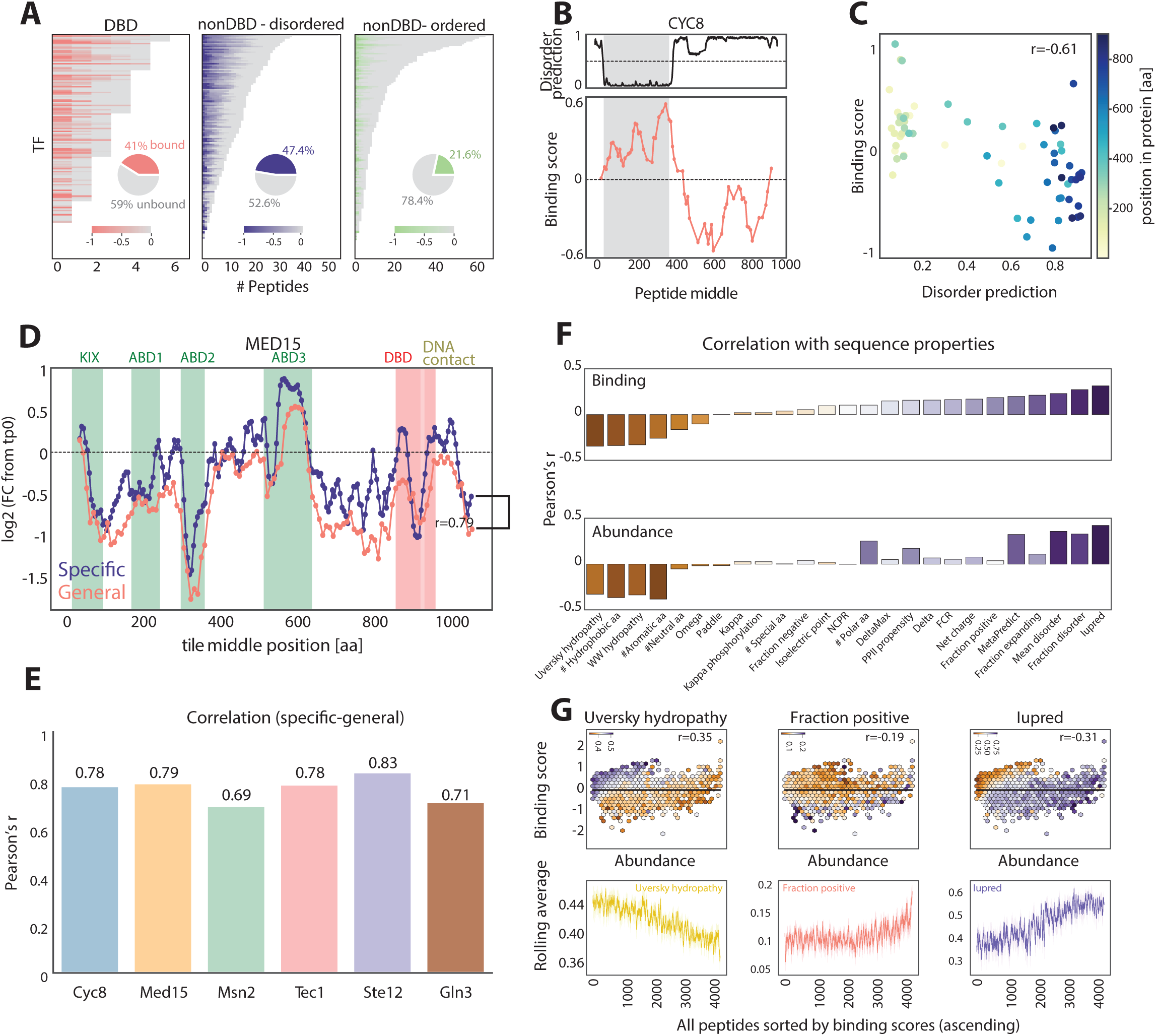
Binding correlates with disorder and charge, and is interrupted by hydrophobic residues. **(A-C)** *Strong binders are enriched in disordered nonDBD peptides:* Peptides in from each TF were classified as those overlapping with the DBD **(A, left)**, disordered (mean IUpred^56^ >0.5) non-DBD **(A, middle)**, and ordered (mean IUpred^56^ < 0.5) non-DBD (**A, right**). The binding score of each TF peptide is shown by color. TFs in each group were ordered by the number of peptides. Pie charts quantify the percentage of peptides with binding score lower than regression slope in each category. **(B-C)** Cyc8, a general co-repressor, is shown as an example: The disorder tendency of Cyc8 is shown **(B, top)** along with the binding scores of each of its composing peptides **(B, bottom)**. A scatter plot comparing the binding score of each peptide with its mean disorder tendency is shown in **(C)**, with the color showing the starting AA position of the peptide. **(D-E)** *High correspondence between general and specific TF libraries:* Specific tiling libraries were generated for Cyc8, Med15, Msn2, Tec1, Ste12, and Gln3, and experiments were repeated for each library. **(D)** The log2 fold change from time point 0 for all Med15 peptides from the general (red) and specific (blue) tiling libraries is shown. Known domain regions^69,70^ are highlighted. **(E)** The correlation between the log2 fold changes of all peptides (comparing between the most similarly positioned peptide) of the indicated TFs in both libraries is shown at the bottom. **(F-G)** *Binding scores are correlated with disorder prediction and negatively correlated with hydrophobic content:* **(F)** The binding scores and abundance of the library peptides were correlated with various protein parameters^51,56,65^ and are presented as bar charts. Disorder tendency and positive amino acid fraction positively correlate with binding scores, while a high fraction of hydrophobic residues disrupts binding. This is further illustrated by the hexagon plots in **(G, top)**, where each hexagon represents the average hydropathy **(left)**, positive fraction **(middle)**, and disorder tendency **(right)** of all peptides with similar abundance (x-axis) and binding score (y-axis). **(G, bottom)** All peptides were sorted by binding scores, and a rolling average was calculated for each attribute shown.

Folded domains may be more sensitive to the precise definition of their boundaries as compared to disordered peptides. To test whether this sensitivity explains the limited binding scores of these domains, we designed new libraries with increased scanning resolution for six tested TFs and coactivators (Msn2, Cyc8, Med15, Tec1, Ste12 and Gln3). In all six cases, the high-resolution profiles correlated well with the low-resolution ones, and corresponded to known domains (**Figure 4D, E**). We conclude that the reduced binding scores of peptides from folded domains, relative to disordered ones is unlikely to result from boundary effects.

Finally, we searched systematically for additional sequence features that correlate with peptide binding scores^51^ (**Figure 4F**). Consistent with the results of the domain library, peptide abundance correlated with low hydrophobicity and high disordered propensity while the (abundance-normalized) binding scores correlated most with the disordered tendency. This correlation of binding scores with the predicted peptide disorder exceeded, in fact, the apparent correlation with peptide charge. Charge effect was observed when restricting the analysis to only disordered peptides (IUpred^56^ >0.5), indicated, for example, by higher fraction of the positive lysine residues, and lower proportion of the polar serine residues within the 100 top scoring ones, as compared to the worst scoring ones (**Figure S4B,C**). Overall, our analysis suggests that both DBDs and disordered nonDBDs contribute to TF binding and provide a detailed map of binding determinants across all TFs.

### Mutation screen of the Msn2 TF

Our tiling libraries pointed at peptides that localized to the tested DNA, often suggesting the presence of multiple such peptides within individual TFs. To further define the sequence features governing this DNA association, we focused on the high-resolution tiling library of Msn2. We measured peptide abundance to define binding scores and selected the top-scoring non-DBD peptide for mutation analysis (residue 209-269, **Figure 5A**). Of note, both abundance and binding scores were well correlated between the low resolution and the Msn2-specific libraries (**Figure S5A**). This peptide contained a high fraction of polar (STNQ) and hydrophobic (LIVM) residues, and a low amount of positively charged residues (KR, **Figure 5B**). Our mutant library was designed for testing different models and included mutations that influenced AA composition (swapping residues between different types of amino acids, e.g., charged KR to hydrophobic LIVM), as well as mutations retaining the overall sequence composition but altering the ordering of residues within the sequence (scrambling residue blocks, clustering of an AA group or randomly distributing same-type residues). Overall, this library included 823 peptides, five repeats of the original peptide, and 15 control peptides encoding for structural regions of enzyme genes not expected to bind DNA. As above, we measured both the binding signal as well as the abundance of each peptide and calculated abundance-normalized binding scores according to the regression line, with both WT and control peptides given the expected binding signals (**Figure 5C, S5B**).

**Figure 5.**
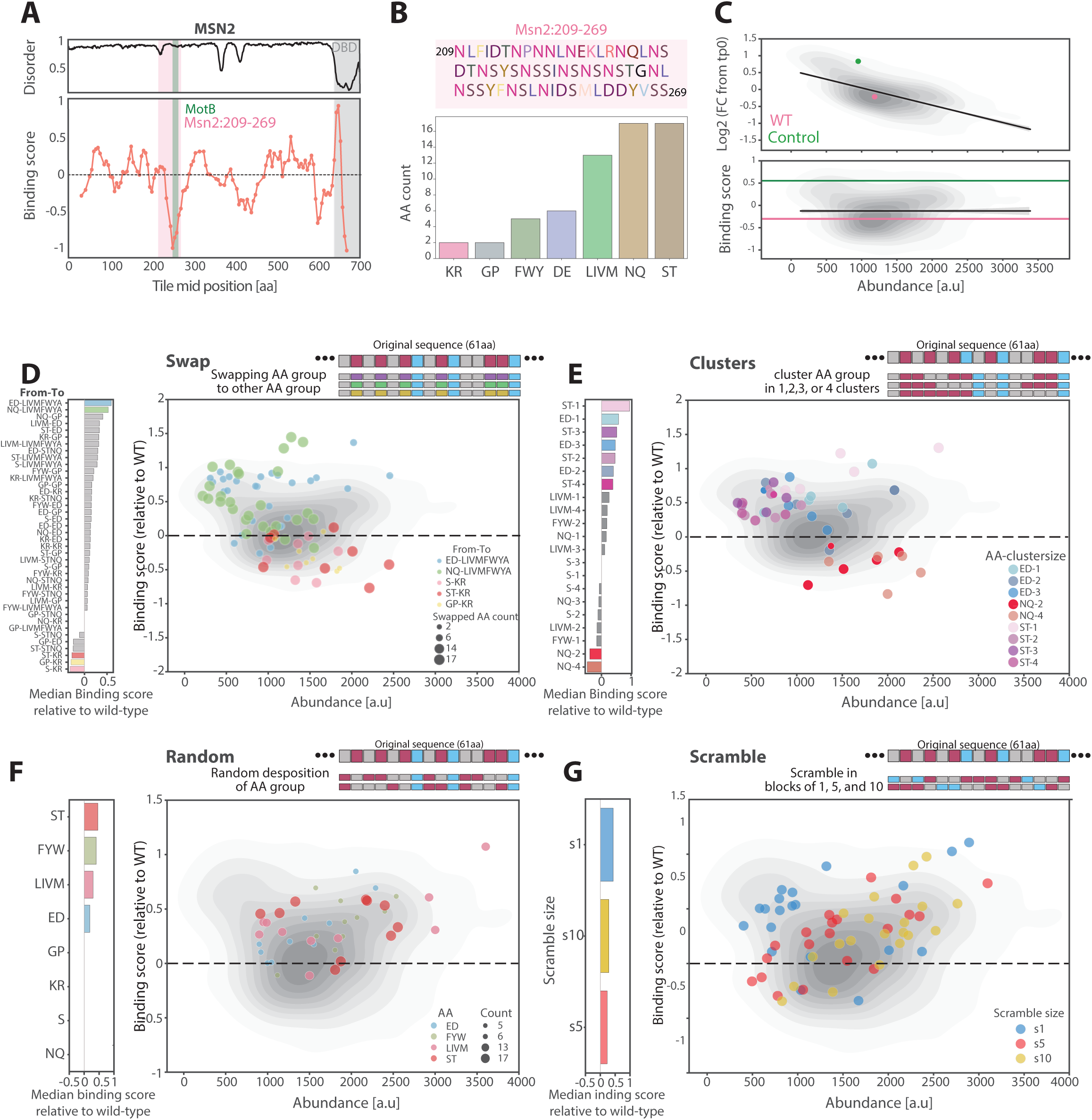
Mutational Analysis of a Disordered Msn2 Peptide Reveals Important Roles of Charged and Polar Residues for Binding. **(A-B)** *Mutational screen of an Msn2 disordered peptide:* **(A)** The Msn2 predicted disorder based on Metapredict^65^ is shown on top, and the binding score of each tile from the specific Msn2 library displayed at the bottom. A disordered peptide showing strong binding, located within residues 209-269 was chosen for the mutational analysis, and its sequence and amino acid composition is shown in **(B)**. **(C)** Over 800 mutation variants were designed (methods), integrated into multiple plasmids containing different target promoters, and binding was measured as described above. Shown in contour density is the log2 fold-change from time zero of all library peptides as a function of abundance. Data rotation by the regression slope was performed as described above. The mean value of the original sequence (WT) is shown in pink, and the mean value of the control peptides is shown in green. **(D-G)** *While most mutations have no effect, clustering and swapping charged residues interrupts binding:* The median binding score relative to the WT sequence of each mutation group is presented on the left as bars for the different mutation types, with extreme groups, showing strong or weak binding, highlighted by color. The binding scores of all peptides included in each of the extreme groups are shown on the right as a function of abundance.

The Msn2 region we selected contains a short 13aa sequence (termed MotB) that interacts with the Med15 coactivator^57^ (**Figure 5A)**. Our library included 470 peptides that were mutated in this sequence, allowing us to test whether this motif is essential for the peptide DNA association we observe. Notably, while MotB-perturbing mutant peptides tended to reduce binding score, the effect size was small with multiple such mutants retaining considerable binding (**Figure S5C**). We conclude that the MotB region stabilizes DNA association, likely through Med15 binding, but is not sufficient to explain the high binding score of the tested Msn2 region.

In agreement with the tiling and domain libraries, increasing the fraction of hydrophobic residues decreased both the peptide abundance and the (abundance-normalized) binding score, as can be seen when replacing the 17 polar (NQ) residues with hydrophobic ones (LIVMFWYA) or when replacing the six negatively charged residues (DE) by hydrophobic ones (**Figure 5D**). Also here, swapping that increased peptide charge led to higher-than-WT binding scores as exemplified by the swapping GP or ST to KR.

Examining next the composition-invariant mutations, pointed at the ED and ST residues whose random deposition or clustering suppressed binding score (**Figure 5E, F**). By contrast, binding scores were largely invariant to the precise positioning of the 17 NQ residues, as neither their random deposition within the sequence nor their clustering was of a significant effect. Finally, scrambling the sequence in three different block sizes reduced binding scores, with a block size of 1 AA having the largest effect, suggesting that both AA composition and distribution affect the binding capacity of this region (**Figure 5G**). Overall, our mutation experiment supports the contribution of positive residues to binding and the inhibitory effect of hydrophobic residues, and further reveals a specific role for the positioning of acidic residues within the tested region of Msn2.

## Discussion

In this work, we introduced pMPBA, a novel method for massive parallel analysis of thousands of peptide-DNA bindings inside cells using sequencing. We applied this method for scanning the sequence space spanned by budding yeast TFs for binding long, and therefore largely non-specific DNA sequence. In this scan, we first tested experimentally the DNA binding capacity of the full set of computationally annotated DBDs. We found large variations both in their stability and in the DNA association of predicted DBDs, at least when expressed independently of the full proteins. It is notable that the apparent strengths of the DBD-DNA association measured in our assays were independent of their motif preferences, consistent with our focus on binding to long and non-specific DNA but were partially explained by DBD charge with positive peptides showing, on average, higher binding scores.

More interesting for us were TF sequences outside known DBDs, where regions influencing genome localization are difficult to detect using existing computational tools. Perhaps unexpectedly, we found that a considerable fraction of peptides lacking DBD annotations showed high binding scores that were comparable, or even exceeded those of known DBDs. Thus, amongst the 100 top-binding peptides, only 17 were annotated as DBDs. Notably, peptides of high DNA binding scores were highly enriched with disordered annotations, suggesting that they lack a stable fold. This association of peptide disorder with DNA was further confirmed within a mutant library, in which >800 mutations of a DNA-associated peptide derived from the Msn2 TF were designed and tested using our system.

Our current design has several limitations. First, and perhaps most importantly, to allow diverse binding modes, we tested peptide binding to a large >2kb DNA segment that included regions coding not only for promoters but also to highly expressed transcripts. Accordingly, the DNA binding signal we detected was largely non-specific, which likely explains the correlation of binding scores with peptide charge and the independence of binding scores from the tested promoter region (∼700bp). In this, our study differs from assays of genomic profiling, including the ChEC-seq method^58^ we previously used to analyze IDR-directed binding^30–33,59^, which report on the preferences for binding at different genomic regions but lacks information on the absolute binding levels. Adapting our method for testing binding to a specific region would require the use of shorter, and a better-defined DNA segment. We also noted that the tested peptides varied significantly in their abundance, which confounds the interpretations and requires normalization of the measured binding scores. Fusing the MNase in the C’ terminus of the tested peptides could potentially protect peptides for the differential N-terminal degradation^60^ to buffer differential abundance, and this will be explored in future studies. Finally, our assay cannot distinguish between direct DNA binding and indirect genomic association and while the former is expected for DBDs, the nonDBD binding showing high binding scores could interact with DNA-localized proteins, including specific TFs or components of the general transcription machinery including the RNA polymerase.

Nonetheless, pMPBA allows, for the first time, a parallel quantification of the association of a library of peptides to a DNA sequence of interest within cells. Using this novel method, we generated the first detailed map of yeast TF binding determinants. With future studies and system modifications, pMPBA will enable us to uncover the molecular grammar underlying TF non-DBD-derived binding specificity.

## Acknowledgments

We thank Gilad Yaakov, Felix Jonas, Dana Bar Zvi, Miri Carmi, and the rest of the Barkai lab members for many fruitful discussions and suggestions. We thank Tally Rosenberg Haimovich, Dalit Peer and Gitit Levi-Cohen from the flow cytometry unit in the Weizmann Institute for their help with cell sorting.

## Materials and Methods

### Strains

All experiments were done on BY4741 strain of genotype MATa his3-Δ1 leu2-Δ0 lys2-Δ0 met15-Δ0 ura3-Δ0.

### pMPBA - Library design and experimental setup

#### Plasmids

plasmids used in this work were based on pRS316 as reported in^42^, were modified to include MNase followed by E2A intein and URA3 selection marker. Plasmid sequence can be found in **Supplementary Table S5**.

#### Domain library

For all budding yeast TFs (not including Zinc cluster), all domain sequences from SGD^61^ were taken. This list of domains was then filtered to include only domains from pfam^45^, SUPERFAMILY^47^, and SMART^46^. The minimal domain size was set to 192 nucleotides (64 AA) and was flanked by sequences necessary for library cloning. The sequences were then modified to remove position the contain restriction sites for the enzymes SgsI and CsiI, that were used for library cloning. Library sequences are found in **Supplementary Table S6**.

#### General TF Tiling library

For each TF, we tiled over the sequence in tiles of 192 nucleotides, with jumps of 33. The last tile of each TF was modified so it will be of 192 nucleotides as well. For Cyc8, Tup1 and Med15, tiles were performed with jumps of 27 nucleotides. Sequences for library amplification and ligation were then added to the sequence. The sequences were then modified to remove position the contain restriction sites for the enzymes SgsI and CsiI. Library sequences are found in **Supplementary Table S6**.

#### Msn2 209:269 mutation library

For the swapping mutations, we first defined 8 AA groups based on similar properties: KR, DE, LIVM, FWY, S, ST, GP, and NQ. All amino acids of a given group were swapped together to all of the following possibilities: L, I, V, M, N, Q, K, R, D, E, S, T, G, P, F, Y, A. The swapping design was split into 4: swapping of all, only the right half, only the left half, or alternating (0-1-0-1) of a given group. The choosing of swapped codons was based on the prevalence of codons in the yeast genome. For all other mutation types the original codons were used. The scramble mutations were designed by taking the original sequence and randomly scrambling it 20 times in blocks of 1,5, and 10 AA. The random mutations were designed by taking each of the 8 AA groups (as in the swapping design) and randomly positioning them, 10 times per group. For the clustering mutation design, we took all residues of each the following AA groups: ‘ED’,’FWY’,’LIVM’,’NQ’,’S’, and ‘ST’, and clustered them in 1,2,3, or 4 locations.

#### Library target promoters

For plasmid target region selection, promoters of the genes GDH1, GDH2, HSP12, HOR7, HXK1, HXT2, HXT7, PCL2, and YAP6 were amplified from genomic DNA and integrated into the target plasmid using the restriction enzymes XmaJI (Thermo Fisher FD1564) on the 5’ end and SfaAI /NhEI (Thermo Fisher, FD2094 and FD0974 respectively) on the 3’ end. Promoter sequences are provided in **Supplementary Table S5**.

### Library cloning and transformation

Libraries were ordered from IDT (4 nmole Ultramer™ DNA Oligos; **Supplementary Table S6**) or Agilent (10 pmol G7220A; **Supplementary Table S6**).

### Library amplification

Libraries were amplified using Herculase II Fusion DNA Polymerase (Agilent, 600677; **Supplementary Table S7** primers 1-7). IDT libraries amplification was done as previously reported^42^. Agilent G7220A libraries were resuspended in 100 ul of Tris pH 8, and 1 ul were used as template for the PCR reaction. Amplification was done using Herculase II Fusion DNA Polymerase and was performed according to the recommendation for 20 cycles. PCR products were separated by electrophoresis on a 1% agarose gel, then libraries were purified from the gel (RBC Bioscience, YDF100) and concentrations were measured using nanodrop machine.

### Digestion and Ligation

Plasmids and libraries were digested using the restriction enzymes SgsI and CsiI (Thermofisher, FD1894 and FD2114 respectively) and ligated together using the fast-link DNA digestion kit (Biosearch technologies, LK0750h) in a respective ratio of 2:1 as previously described^42^.

### Bacterial transformation

Ligation products were then transformed into E. cloni® Electrocompetent Cells (Biosearch technologies, LC601172). Transformation was done according to the provided protocol, at 4°C to increase yield. Cells were evenly plated on ten 14 cm LB-amp plated and grown over night at 37°C. A small amount of cells was plated for counting the total number of transformants (5-30M for transformants for each library) and 10-20 colonies from each transformation were collected for library insertion validation using PCR (**Supplementary Table S7**, primers 8-9).

### Plasmid Extraction

Colonies growing on the 14 cm plates were scraped and washed using LB media and collected in 250 ml bottles. Plasmids were extracted using the NucleoBond Xtra Maxi kit (Machery-Nagel, 740414.10), according to the provided protocol, yielding 1000-1500 ng/ul.

### Yeast transformation

To transform yeast with plasmids carrying the library the BY4741 strain was transformed using the LiAc/SS DNA/PEG method as previously reported^42^. Following transformation cells were inoculated in fresh 25 ml SD -URA and were grown to stationary phase for 72 hours (shaking at 30°C).

### Experimental procedure

Experimental procedure was similar to the previously reported MPBA procedure^42^. Library-transformed stationary cultures were refreshed using 10 ml SD -URA media (per sample) to reach OD_600_ of 4 after ∼10 cell divisions at 30°C. This 10 ml refreshed sample will be later on splited into two 5 mLl samples that will be subjected to different treatments: The non-activated sample (time point 0) will be taken directly to DNA extraction (see below), and the rest of the sample will be subjected to MNase activation (see below) before DNA extraction.

### MNase activation

This procedure was performed as previously described^42^. In brief, cultures (OD_600_=4) were pelleted at 1500 g and resuspended in 1 mL Buffer A (15 mM Tris pH 7.5, 80 mM KCl, 0.1 mM EGTA, 0.2 mM spermine, 0.5 mM spermidine, 1 × Roche cOmplete EDTA-free mini protease inhibitors, 1 mM PMSF), and then transferred to deep well plate. Cells were washed twice in 500 μL Buffer A, pelleted, and resuspended in 150 μL Buffer A containing 0.1% digitonin. Then, cells were transferred to a 96-well plate (PCR-96-FLT-C, Axygen) for permeabilization (30°C for 5 min). CaCl_2_ was added to a final concentration of 2 mM. Mnase was activated for different times, as specified in the text. Next, 100 μL of stop buffer (400 mM NaCl, 20 mM EDTA, 4 mM EGTA and 1% SDS) were mixed with 100 μL of sample. Proteinase K was then added, and incubated at 55°C for 30 min.

### DNA extraction

Plasmids DNA were extracted both from MNase-activated cells and non-activated cells using the MasterPure Yeast DNA Purification Kit (Lucigen Corporation, MPY80200), with the modification that were previously reported^42^.

### Library preparation

Samples were 1 X SPRI (AMPure XP, A63881) cleaned and resuspended in 30 ul elution buffer (10mM Tris-HCl, pH 8) and then diluted 1:6. 1 ul was used as a template for PCR reaction for amplifying noncleaved variants and sample barcoding (**Supplementary Table S7**, primers F: 10-41, R: 42-47; 28 cycles) using the KAPA HiFi HotStart ReadyMix (Roche, KK2602). PCR products were verified using 1% agarose gel electrophoresis, and 1X SPRI cleaned. DNA concentrations were measured using Qubit™ Flex Fluorometer (Invitrogen), and samples were normalized to equal amounts, then pooled together. 1 ng was taken as a template for a 10 cycles PCR reaction (**Supplementary Table S7**, primers F: 48, R: 49-57) using KAPA HiFi HotStart ReadyMix for adding IIlumina indices and machine adapters. Samples were 0.5X reversed SPRI cleaned, and then 0.8X SPRI cleaned. Then concentrations and library quality were validated using Qubit and tape station (Agilent).

### NGS library preparation and sequencing

Libraries were sequenced using the NovaSeq 6000 machine. Runs were performed with the SP200 kit (20040719), parameters: R1 - 61 cycles, Index1 - 8 cycles, Index2 - 8 cycles, R2 - 61 cycles. In each run, 5% PhiX DNA was added to increase complexity.

### Cell sorting

In order to calculate peptide abundance, peptide libraries were transformed to a plasmid containing a GFP (instead of an MNase) as described above. Plasmid sequence is found in **Supplementary Table S6**. Following transformation, cells were allowed to grow for 2 days with back dilution to eliminate multi-plasmid containing cells. Cells were sorted using BD FACS Aria II. Single cells were sorted to eight bins based on GFP fluorescence. Bins were defined based on the overall GFP distribution, with each bin containing 10% of the population. For each bin, at least two repeats were collected, each containing ∼15000 cells for the small libraries (Domains, Msn2-specific and Msn2 :209-269 mutations) and ∼100,000 cells for the larger general TF tiling library. Following sorting, cells were allowed to grow for 2-3 days following DNA extraction and library preparation and sequencing as described above.

### Analysis and statistics

#### Code and data availability

Sequencing data generated in this study have been deposited in GEO under accession GSE272404. Code and processed data file can be found in https://github.com/sagieb/pMPBA.

#### Peptide counts

Snakefile was constructed using SnakeMake^62^. Raw reads were demultiplexed using bcl2fastq (Illumina), and adaptor were filtered out using cutadapt^63^. For domains, tiling, and TF specific tiling libraries, reads were mapped to a reference genome containing all designed library variants using bowtie2^64^. For Msn2 pep3 mutation library, reads were assigned to each peptide by its unique barcode sequence. Reads were then normalized to one million to account for sequencing depth and samples with either low amount of reads or low correlation between repeats were removed. Technical repeats were averaged. The Fold change of each peptide count from timepoint 0 was calculated. The median fold change values across biological repeats and promoters was then calculated and used for downstream analysis.

#### Abundance calculation

Read count of GFP-fused libraries was calculated as described above, resulting in a normalized read count in each bin for each peptide. For each peptide, the read fraction in each bin was calculated and multiplied by the median GFP signal of the bin. Peptides with low number of reads were set as nan. The resulting values were then summed to calculate the abundance value of each peptide. Peptides with standard deviation larger than 1000 between repeats were removed.

#### Binding score (Abundance normalized binding)

A linear regression line between peptide abundance and fold change (timepoint 360’’ from timepoint zero) was calculated. The distance of each point from the regression line was defined as the normalized binding score. For presentation, centroid rotation by the regression slope was performed to rotate the data along the regression plane.

#### Unique domains (Figure2)

The domain library contained overall 351 domains, including mostly overlapping peptides. Domains belonging to the same TF and vary in up to 20 AA in their starting position were considered overlapping domains. Overlapping domains were filtered so that the domain with the best binding score was kept and the others discarded.

#### Sequence chemical properties (Figure 2 and 5)

localCIDER^51^ was used to calculate most chemical properties. Metapredict^65^ and IUpred2^56^ were used to calculate the mean disorder prediction value of each peptide. Mean disorder propensity was calculated by averaging the disorder propensity of each AA as defined in^66^.

#### Tiling classification with InterPro (Figure 3)

For all examined TFs, all domain listed in InterPro^52^ were taken. Each Tile was assigned to its closest domain, in case at least half of the domain was included within the tile sequence. Domains from Gene3D, Coils, and SignalP databases were removed from this classification.

#### Motif Euclidean distance (Figure S2B)

The Euclidean distances between TF motifs was calculated using the position weight matrix (PWMs) as in^42^.

#### Motif count (Figure S2A)

To count the number of motifs present in each sequence, a consensus motif sequence was taken from^42^ based on the TF PWMs. The number of occurrences present within each promoter sequence was counted.

#### Sequence conservation (Figure S2C)

Conservation scores were calculated by taking multiple sequence alignments (MSA) of each TF from YGOB^67^. For each MSA, Shannon entropy scores were calculated for each position, taking only positions without a gap in the cerevisiae sequence. Evolutionary conservation scores were calculated by taking the Shannon entropy scores (scaled between 0-1) and subtracting from 1, so that higher scores indicate greater conservation. Finaly, evolutionary conservation vectors were trimmed according to peptide position coordinates and the median value was taken as the peptide conservation score.

#### Activation activity prediction (Figure S4A)

PADDLE^38^ was used to calculate activation activity prediction, taking the median value of each sequence.

## Supplemental information

**Supplementary** Figures 1-5

**Table S1**: Domain library data

**Table S2:** General TF tiling library data

**Table S3:** Msn2-specifc tiling library data

**Table S4:** Msn2:209-269 mutation library data

**Table S5:** Plasmids and promoters sequences

**Table S6:** Libraries sequences

**Table S7:** Primer list

---

**Figure S1.**
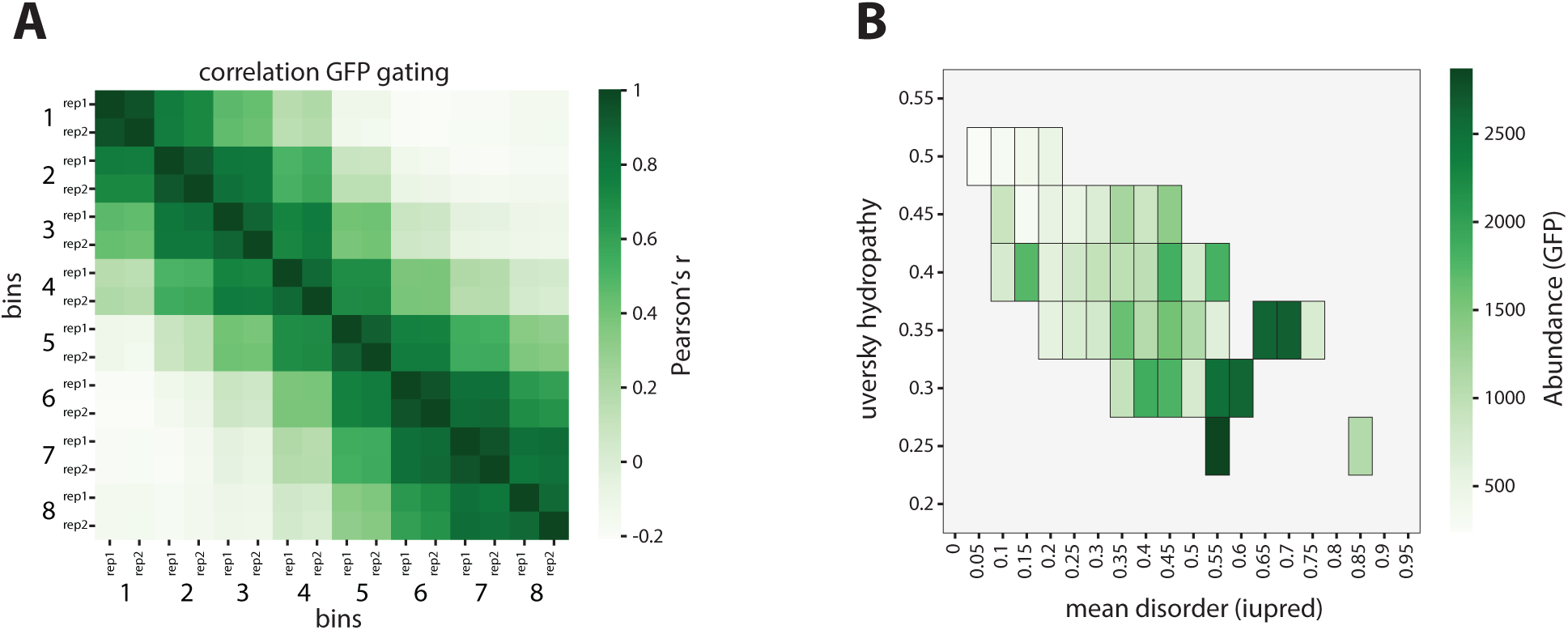
Peptide abundance is affected by disorder propensity and hydropathy. **(A)** *Similarity of domain abundance between repeats and adjacent bins:* The cells were sorted into eight bins based on GFP fluorescence. Shown are the correlations of domain abundances along the bins and between technical repeats. **(B)** *Correlation between predicted disorder, hydropathy and abundance:* Shown is the domain disorder prediction (IUpred^56^) as a function of hydropathy. The average abundance levels of domains for which the disorder and hydropathy measures are found within a given rectangle are shown in color.

**Figure S2.**
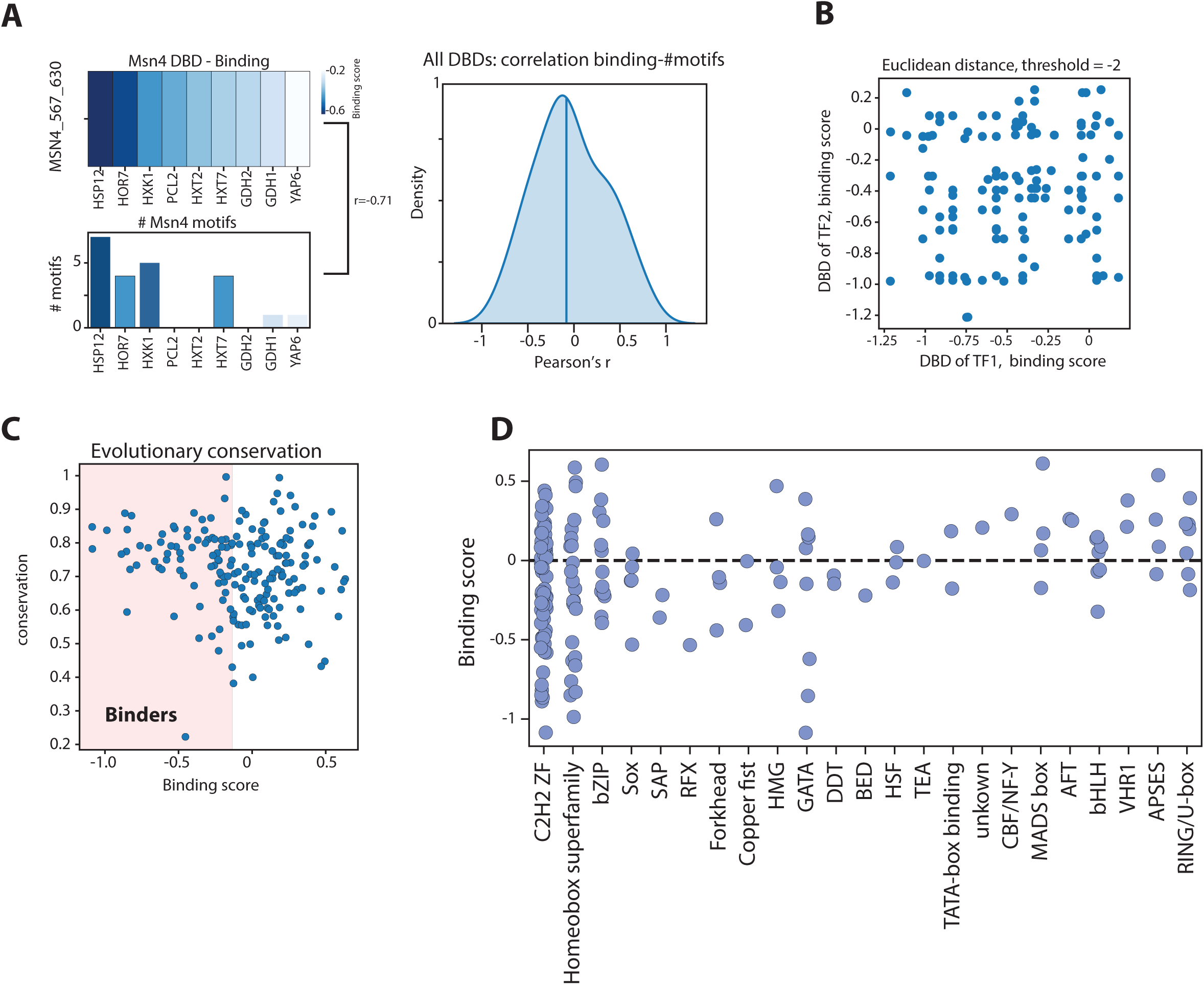
High conservation of strong binders and low correspondence between motif presence and binding strength. **(A)** *Motif abundance poorly correlates with binding:* Shown on the left is binding score of the Msn4 DBD on each tested promoter (top) and the number of preferred motifs (AGGGG) found within each promoter sequence (down). Shown on the right is a distribution of the correlations between binding scores of each TF domain on each promoter and the number of preferred motifs found in the sequence. **(B)** *Strong binders are evolutionary conserved:* Shown are the domain binding scores as a function of evolutionary conservation (methods). **(C)** *Similar DBDs show differential binding:* Shown are the binding scores of all DBD pairs for which the Euclidean distance between the PWMs was higher than −2 (methods). Ohnologs, shown in Figure 2G, were excluded from this analysis. **(D)** *Binding by DBD family:* Shown are the binding scores of all measured peptide domains separated by TF family.

**Figure S3.**
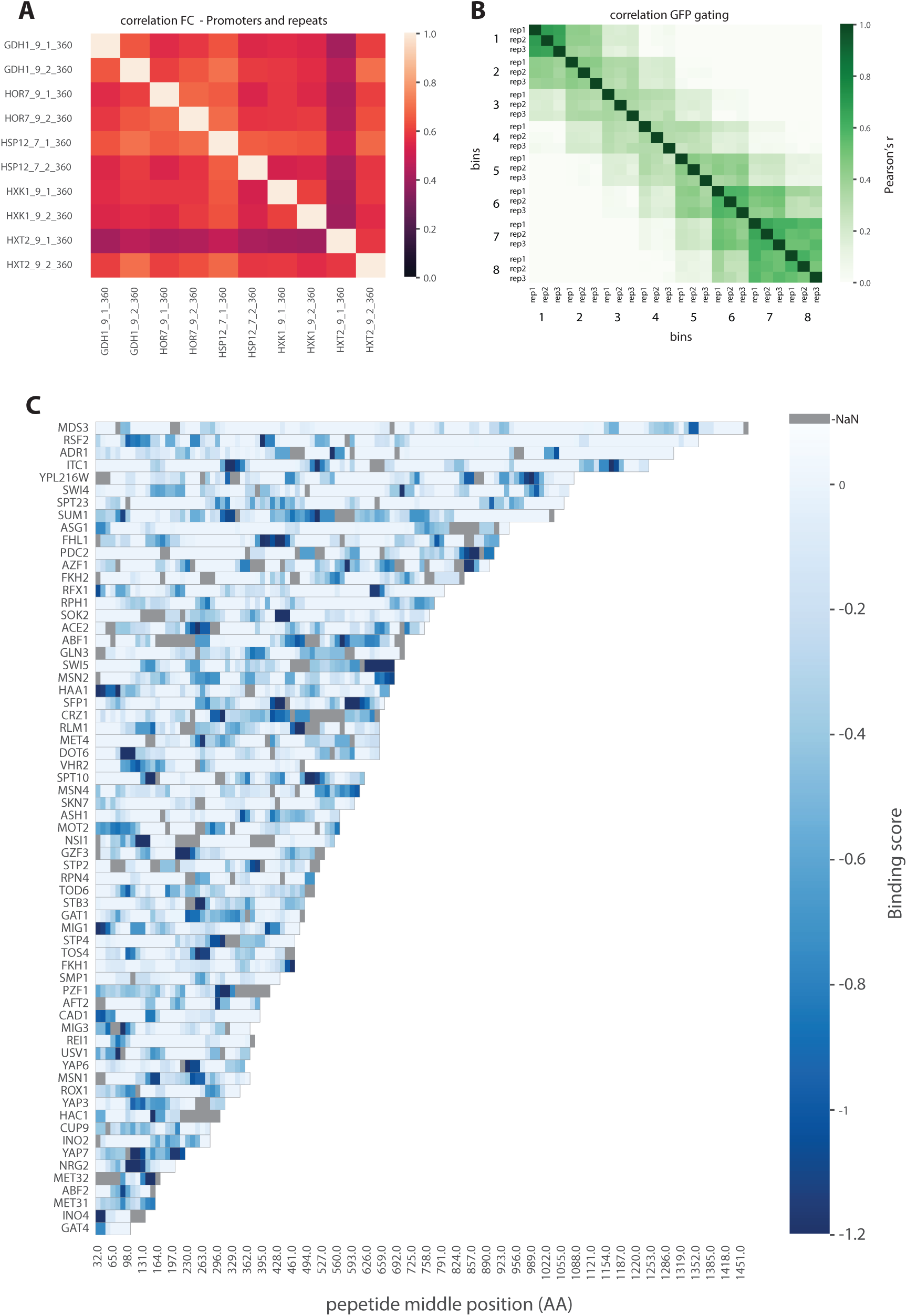
Correlation between repeats, promoters, and abundance of proximal GFP bins. **(A)** *High binding similarity between promoters:* Shown is the correlation of log2 fold-changes of all peptides of the tiling library measured across different promoters. Two biological repeats of each promoter are presented. **(B)** *Similarity of abundance between repeats and adjacent bins:* The cells were sorted into eight bins based on GFP fluorescence. Shown are the correlations of domain abundances along the bins and between technical repeats. **(C)** *Peptide binding map:* Shown are the binding scores of all tiles of TFs with at least one peptide in the top 100 binders of the TF tiling library as in Figure 3E. TFs are ordered based on sequence length.

**Figure S4.**
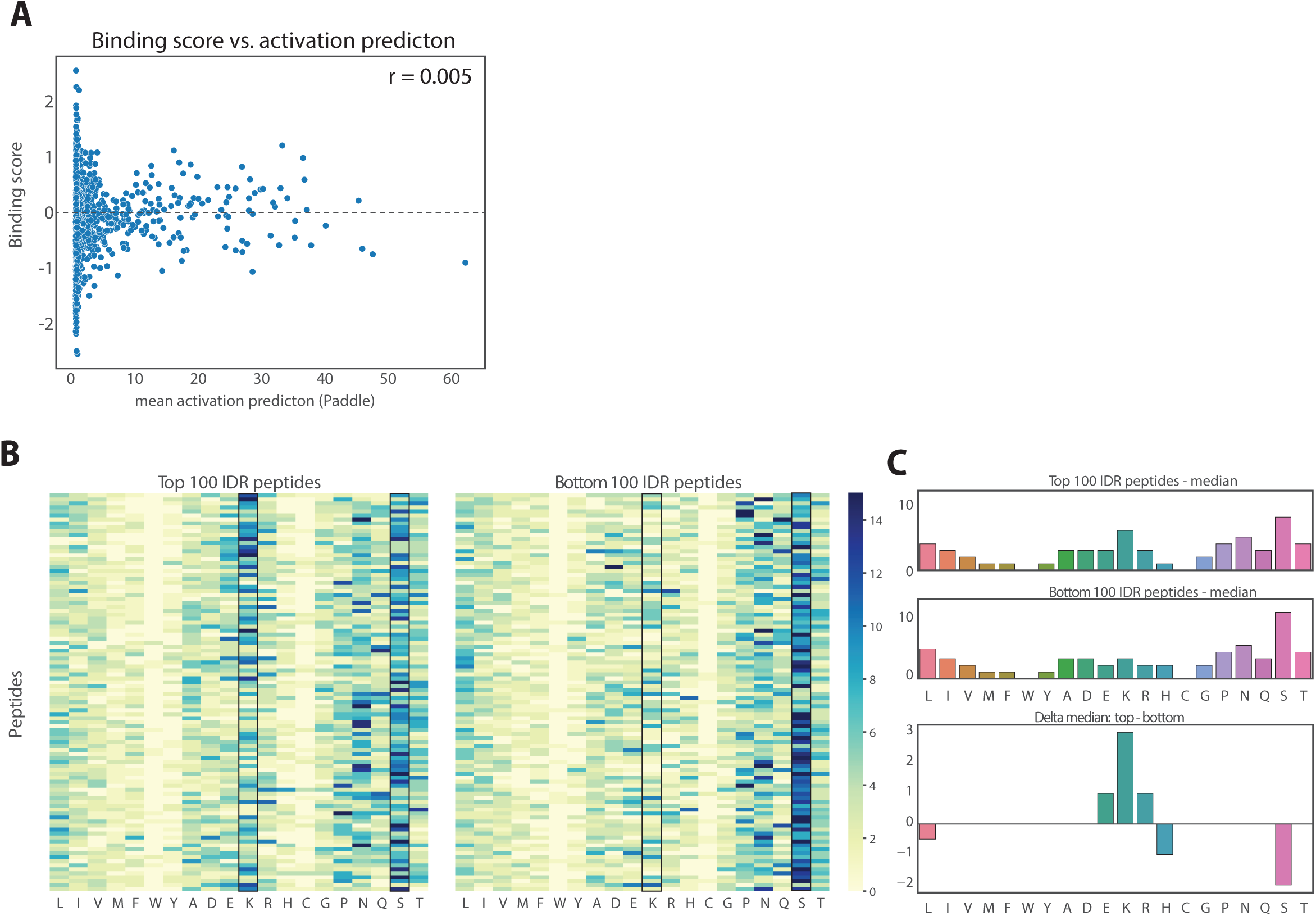
Distinct IDR properties of weak and strong binders. **(A)** *Binding score poorly corresponds to peptide activation capacity*: Shown are the abundance-normalized binding score as a function of the mean predicted activation capacity (calculated by PADDLE^38^) of each library peptide. **(B-C)** *Charged amino acids are enriched in strong IDR binders*: Shown on the left is the distribution of amino acids of each of the top 100 IDR binders in the tiling library (IUPred ^56^score >0.5), and on the right the same presentation for the bottom 100 IDR binders (IUpred^56^ score < 0.5). The median number of each amino acid of the top (right, top) and bottom (right, middle) IDR binders are also shown, together with the delta between these medians (right, bottom).

**Figure S5.**
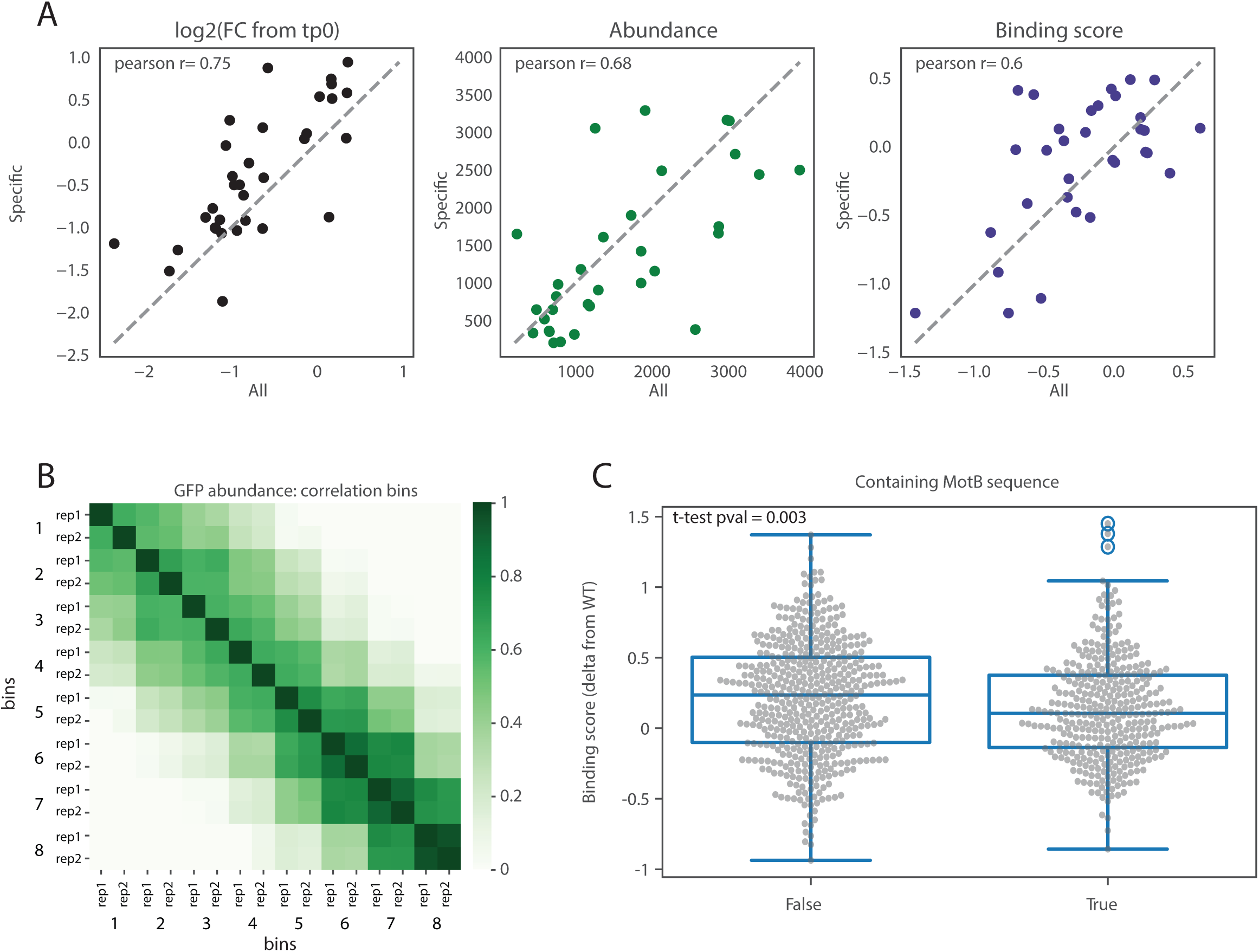
Quality controls of the specific Msn2 and peptide 209-269 mutation libraries. **(A)** *High correspondence of Msn2 peptides between different libraries*: A comparison of Msn2 peptides between the specific Msn2 tiling library (129 peptides in total) and the general TF tiling library (7111 peptides in total). Shown is the log2 fold change from time point (left), abundance (measured by GFP, middle), and abundance-normalized binding score (right) for each Msn2 peptide in each library. Note that due to difference in peptide annotations between the libraries, each peptide from the Msn2 specific library is compared to the most similar peptide of the general TF tiling library. **(B)** *Similarity of in abundance between repeats and adjacent bins in the mutation library of Msn2 peptide 209-269:* The cells were sorted into eight bins based on GFP fluorescence. Shown are the correlations of domain abundances along the bins and between technical repeats. **(C)** *Small effect of previously described activation determinant found within Msn2 peptide 209-269:* Shown are the distributions of binding scores of peptides where the previously defined activation determinant^57^ is mutated (left) or intact (right). While there is a significant contribution of this determinant to the total binding score (p. value = 0.003), the effect size (Cohen’s D = 0.2) is low.

